# Retinoic acid generates a beneficial microenvironment for liver progenitor cell activation in acute liver failure

**DOI:** 10.1101/2024.01.23.576749

**Authors:** Sai Wang, Frederik Link, Stefan Munker, Wenjing Wang, Rilu Feng, Roman Liebe, Yujia Li, Ye Yao, Hui Liu, Chen Shao, Matthias P. A. Ebert, Huiguo Ding, Steven Dooley, Hong-Lei Weng, Shan-Shan Wang

**Affiliations:** Department of Medicine II, University Medical Center Mannheim, Medical Faculty Mannheim, Heidelberg University, Mannheim, Germany; Department of Medicine II, University Hospital, LMU Munich, Munich, Germany; Liver Center Munich, University Hospital, LMU; Munich, Germany; Beijing Institute of Hepatology, Beijing You’an Hospital, Capital Medical University, Beijing, China; Department of Endocrinology and Metabolism, Renji Hospital, School of Medicine, Shanghai Jiao Tong University, Shanghai, China; Clinic of Gastroenterology, Hepatology and Infectious Diseases, Otto-von-Guericke-University, Magdeburg, Germany; Department of Pathology, Beijing You’an Hospital, Affiliated with Capital Medical University, Beijing, China; Molecular Medicine Partnership Unit, European Molecular Biology Laboratory, Heidelberg, Germany; DKFZ-Hector Cancer Institute at the University Medical Center, Mannheim, Germany; Department of Gastroenterology and Hepatology, Beijing You’an Hospital, Affiliated with Capital Medical University, Beijing, China

**Author notes:** Corresponding author: Shanshan Wang Beijing Institute of Hepatology, Beijing You’an Hospital, Capital Medical University, Beijing, China No.8 Xi Tou Tiao, You An Men Wai, Feng Tai District,Beijing, 100069, China. These authors contributed equally to this work (Sai Wang, Frederik Link).

**Keywords:** HSC activation, retinoic acid, RARα, LPC, ALF

## Abstract

**Objective:** When massive necrosis occurs in acute liver failure (ALF), rapid expansion of hepatic stem cells called liver progenitor cells (LPC) in a process called ductular reaction (DR) is required for survival. The exact underlying mechanisms of this process are not known to date. In ALF, high levels of retinoic acid (RA), a molecule known for its pleiotropic roles in embryonic development, are secreted by activated hepatic stellate cells (HSCs). We hypothesized that RA plays a key role during DR in ALF.

**Methods:** RNA-Seq was performed to identify molecular signaling pathways affected by all-trans retinoid acid (atRA) treatment in HepaRG LPC cells. Functional assays for RA were performed in HepaRG cells with atRA treatment as well as co-culture with LX-2 cells *in vitro*, and liver tissue of patients suffering from ALF *in vivo*.

**Results:** Under ALF conditions, activated HSCs secreted RA, inducing RARα nuclear translocation in LPCs. RNA-seq data and investigations in HepaRG cells revealed that atRA treatment activated the WNT-β-Catenin pathway, enhanced stemness genes (SOX9, AFP, et.al), promoted energy storage, and elevated the expression of ATP-binding cassette (ABC) transporters depending on RARα nuclear translocation. Further, atRA treatment-induced pathways were confirmed in a co-culture system of HepaRG with LX-2 cells. Patients with ALF who displayed RARα nuclear translocation in LPC had significantly better MELD scores than those without.

**Conclusion:** In ALF, RA secreted by activated hepatic stellate cells promotes LPC activation, a prerequisite for subsequent LPC-mediated liver regeneration.

## Introduction

Massive hepatic necrosis (MHN) occurs during the terminal and most lethal stages of acute liver failure (ALF) (1), leaving only a few remaining functional hepatocytes(2). Hepatocytes perform most functions of the human liver, for example, the production of bile acids and various plasma proteins including coagulation factors, nutrient storage and metabolism, elimination of xenobiotics, steroid hormones and cytokines to support systemic homeostasis (3, 4). Some patients with ALF in whom most of the hepatocytes are damaged however, can recover following acute decompensation. The underlying mechanisms are thought to be dependent on liver progenitor cells (LPCs) (2, 5, 6). LPCs are stem-like cells that reside along the canals of Hering and biliary ductules of healthy livers. They proliferate rapidly in severely damaged livers and can be identified by immunohistochemical staining for biliary markers, such as CK19 and CK7 (1, 7). The mechanisms that regulate LPC activation, most likely through signals from the surrounding microenvironment, remain largely unknown.

Hepatic stellate cells (HSCs) constitute 5-8% of all liver cells (8) and contain approximately 80% of the body’s vitamin A (retinol). Retinol is metabolized into RA by two steps and has pleiotropic functions in embryonic development (9) and liver homeostasis (10). *In vivo,* during liver injury or *in vitro,* during cell culture, HSCs are activated, secrete retinoids rapidly and upregulate their production of profibrogenic cytokines (e.g., TGF-β1) and extracellular matrix components (11, 12). The retinoids liberated from lipid droplets during HSC activation can be used for retinoic acid formation, whose biologic effects are mainly mediated through two nuclear receptors: retinoic acid receptors (RARs) and retinoid X receptors (RXRs) (13, 14). atRA is believed to be the endogenous ligand for RARs as it activates RARs at nanomolar levels, especially RARα, the most abundant isoform in the liver (15). According to previous studies, atRA-RAR/RXR pathways promote fatty acid oxidation, lipid metabolism whilst inhibiting lipid and bile acid synthesis in hepatocytes (16–18). atRA also demonstrates inhibitory effects on the expression of profibrotic genes, including TGF-β1, COL1A1, MMP2, and α-SMA in HSCs through RAR-dependent down-regulation of myosin light chain 2 (MLC-2) expression on a transcriptional level (19–22). In addition, RA has been shown to reduce hepatic inflammation by repressing the production of pro-inflammatory cytokines in dendritic cells, monocytes, macrophages, and T cells in an RAR/RXR-dependent manner (23–25) thereby regulating immune cell differentiation and activation (23, 26). However, the effects of RA on liver progenitor cells (LPCs), especially during acute liver failure (ALF), remain unclear.

In this study, we investigated the effects of RA on the HepaRG liver progenitor cell line using RNAseq *in vitro* and further verified our findings *in vivo* using liver tissue from ALF patients.

## Material and Methods

### Reagents and antibodies

atRA (cat. #R2625-100MG, Sigma-Aldrich, Taufkirchen, Germany); Human RARα siRNA (cat. #sc-29465, Santa Cruz Biotechnology, Santa Cruz, CA, USA); RARα antibody (cat. #PA5-39870, Cell Signaling Technology, Danvers, MA, USA); Active β-Catenin antibody (cat. #05-665, Millipore, Darmstadt, Germany); β-Catenin antibody (cat. #sc-7963, Santa Cruz Biotechnology); SOX9 antibody (cat. #82630, Cell Signaling Technology); AFP antibody (cat. #HPA010607, Sigma-Aldrich); MRP1 antibody (cat. #sc-18835, Santa Cruz Biotechnology); Alexa Fluor 555 goat anti-mouse IgG (cat. #A-21422, Invitrogen, Waltham, MA, USA); Alexa Flour 488 goat anti-rabbit IgG (cat. #A-11008, Invitrogen); DRAQ5 (cat. #4084L, Cell Signaling Technology); GAPDH antibody (cat. #32233, Santa Cruz Biotechnology); BODIPY (D-3922, Life Technologies, Waltham, MA). Insulin-Transferrin-Selenium (ITS -G) (100X) (cat. #41400045, Thermo Fisher Scientific, Waltham, MA).

### Liver tissue specimens of patients

Twenty-one liver tissues of patients with ALF were collected from Beijing You’an Hospital, Affiliated with Capital Medical University and Department of Medicine II, University Medical Centre Mannheim, Medical Faculty Mannheim, Heidelberg University. In this study, ALF was defined as an acute decompensation occurring in individuals regardless of the presence of prior liver damage. Acute decompensation was diagnosed as: (i) Total bilirubin (TBIL) > 10 ULN or increased TBIL 1mg/d with or without grade 2 to 3 ascites within <2 weeks; (ii) overt hepatic encephalopathy; (iii) gastrointestinal haemorrhage; (iv) International normalized ratio (INR) ≥1.5, (v) bacterial infections (spontaneous bacterial peritonitis, spontaneous bacteraemia, urinary tract infection, pneumonia, cellulitis) (27). The liver tissue from 21 ALF patients was collected when they underwent liver transplantation. The study protocol was approved by the local ethics committees (Jing-2015-084 and 2017-584N-MA). Written informed consent was obtained from patients or their representatives. For liver tissues collected by liver transplantation, allocation and timing of LTx were governed by the China Liver Transplant Registry (CLTR) (28), an official organization for scientific registry authorized by the Chinese Health Ministry, according to the Model for End-Stage Liver Disease (MELD) score of the patient.

### Primary human stellate cell isolation

Primary human HSCs (phHSCs) were isolated by the Cell Isolation Core Facility of the Biobank Großhadern, University Hospital, LMU Munich, using discontinuous density centrifugation with percoll (29). phHSCs were grown in 10% DMEM supplemented with 1% L-glutamine, 1% P/S and 10% FBS. All cells were kept in a humidified 37°C incubator enriched with 5% CO_2_.

### Cell culture and treatment

HepaRG cell line is a human bipotent liver progenitor cell line (30). HepaRG cells were kept in William’s E medium supplemented with 10% FBS, 1% L-glutamine, 1% P/S, 50µM hydrocortisone hemisuccinate, and 5µg/ml insulin. LX-2 HSCs were kept in DMEM supplemented with 2% FBS, 1% L-glutamine, and 1% P/S. Following overnight culture, HepaRG cells were transfected with control or RARα siRNA, treated with control or 10µM atRA for 24, or co-cultured with LX-2 HSCs in trans-well for 72h before the sample collection for further analysis. For inducing HepaRG differentiation to hepatocyte, the cells were cultured in differentiation medium (William’s E medium supplemented with 10% FBS, 1% L-glutamine, 1% P/S, 1x Insulin-Transferrin-Selenium (ITS -G) (100X), 50ng/ml EGF, 40ng/ml dexamethasone, and 10mM nicotinamide) for 2, 4, or 7 days.

### MTT assay

Proliferation measurements were performed using the thiazolyl blue tetrazolium bromide (MTT) assay. 4,000 cells per well were seeded in quadruplicate in a 96-well plate. After attachment, cells were treated as described in the section *cell culture and treatment*. At the end of all other experiments, the remaining cell culture medium was discarded, and 100 µl of new culture medium and 10 µl MTT reagent was added to each well and incubated for 4 h at 37°C to enable the formation of formazan crystals. After aspiration of the medium, 100 µl/well DMSO were added, and the plate was incubated on a shaker at room temperature for 10 min to dissolve the formazan crystals. Absorbance was measured at 570 nm using the Infinite 200 Spectrophotometer.

### Western blotting

For whole-cell lysates, cultured cells were washed with ice-cold PBS and dissolved in RIPA lysis buffer (1% Triton X-100, 50 mMTris [pH 7.5], 300 mM NaCl, 1 mM EGTA, 1 mM EDTA, and 0.1% SDS), with freshly added phosphatase and protease inhibitors. Protein concentrations were measured using the Bio-Rad protein assay kit according to the manufacturer’s instructions and were quantified by the Tecan Infinite M200 via absorbance measurement at 690 nm. 20 µg proteins were separated by SDS-PAGE (10– 12% gels) and were blotted onto a nitrocellulose membrane (Millipore, Billerica, MA, USA). The membrane was blocked with 5% BSA in TBST at room temperature for 1h. Subsequently, the membrane was incubated with primary antibodies overnight at 4 °C. Next day, the membrane was incubated with HRP-linked anti-mouse or anti-rabbit secondary antibody following PBST washing three times. Signals were visualized by the Supersignal Ultra solution (Pierce, Hamburg, Germany) and recorded by the imaging system Fusion SL4 (PEQLAB, Germany). The target protein bands were quantified by Fluorchem Q system (ProteinSimple, California, USA). For liver tissue proteins, 20 µg extracted proteins were used for Western blotting. Each Western blotting experiment was repeated at least three times.

### Transfection and RNA interference

For gene silencing experiments, cells were transfected by 50 nM siRNA using Lipofectamine RNAiMAX (Invitrogen) following the manufacturer’s protocol. Cells were harvested 48 hours post-transfection.

### RNA isolation and qRT-PCR

RNA was isolated from liver tissues or cultured cells with the TRIZOL reagent (Sigma-Aldrich). 500 ng RNA were used for first-strand cDNA synthesis with a cDNA synthesis kit (Thermo Fisher). A 20 µl mixture containing 5 µl cDNA (diluted 1:10), 4 µl SYBR Green, 10μM of both forward and reverse primer were used for real time PCR by a StepOnePlus Real-Time PCR instrument (Applied Biosystem). PCR amplification cycling conditions comprised a polymerase activation step for 10 minutes at 95 °C, and a subsequent amplification step for 15s at 95 °C and 1 minute at 60 °C for 40 cycles. A melting curve was established to validate the specificity of each PCR analysis. Relative quantification of target genes was normalized against the house keeping gene PPIA. Primer sequences were retrieved from the PrimerBank® (Massachusetts General Hospital, Boston, MA, USA) online resource and ordered from Eurofins Genomics (Eurofins Scientific, Luxemburg, Luxemburg) (Table S1).

### Chromatin Immunoprecipitation

Chromatin immunoprecipitation was performed according to a previous description (31). Approximately 1 × 10^6^ HepaRG cells treated with control or atRA for 24h were collected and prepared for ChIP assay. 2 µg of RARα antibody or rabbit IgG were used for the incubation with the samples. The control or RARα antibody-pulled down DNA was extracted and used for PCR analysis. Primer sequences of *CTNBB1* and *SOX9* for ChIP assay were in Table S1.

### Immunofluorescence (IF) staining

Cultured cells were fixed on slides with 4% PFA for 10 min. After washing with PBS for three times, slides were permeabilized with 0.5% Triton X-100 in PBS for 10 min. After rinsing the samples with PBS, cells were blocked with 0.025% Triton X-100 and 1% BSA in PBS for 1h at room temperature. Subsequently, the slides were incubated with primary antibodies in PBS, 0.025% Triton X-100 and 1% BSA at 4 °C overnight. The next day, all slides were incubated with fluorochrome-conjugated secondary antibodies and DRAQ5 (Invitrogen) for 1h followed by three washes with PBS plus 0.025% Triton X-100. Images were scanned under a Leica Confocal TCS SPE.

### Immunohistochemistry staining (IHC)

Liver sections of 4 µm thickness were used for staining. After deparaffinization, sections were washed in PBS for 3 times. Heat-induced antigen retrieval was performed in EDTA buffer and endogenous peroxidase activity was blocked by Dako blocking reagent. The slides were incubated with primary antibody or PBS (negative control) overnight at 4°C cold room. Next day, the slides were incubated with streptavidin-conjugated horseradish peroxidase antibody at room temperature for 1 hour. Staining was visualized with diaminobenzidine (DAB) and samples were counterstained with hematoxylin. The details of the protocal have been described previously (32).

Stained slides were scanned using a whole slide scannin microscope (Aperio Scanscope CS, Leica Biosystems, Nussloch GmbH, Germany). A pathologist who was blinded as to the patients’ characteristics evaluated every tissue section independently.

### Lipid droplet staining

AML12 cells or cryosections (4µm thick) were washed briefly with PBS for 3 times, followed by incubation with BODIPY (1:250) in PBS for 30 minutes at room temperature. After washing with PBS twice, the cells or sections were fixed with 4% PFA and stained with phalloidin and DRAQ5, and analysed by confocal microscopy as above.

### Time–of–flight mass cytometry

Time–of–flight mass cytometry was performed for analysis of expression of the 26 related proteins in atRA treated HepaRG cells as previously described (33). Briefly, HepaRG cells were harvested and labelled with Cisplatin to distinguish the dead cells. Then, the cells stained with antibody cocktail against target markers and DNA intercalator. Finally, HepaRG cells was detected on the Helios mass cytometer. Based on normalized protein expression of 26 markers (Table S2), t-distributed stochastic neighbour embedding (t-SNE) algorithm methods was used to analyse the collected data after the normalized and cleaned via Cytobank (https://www.cytobank.org/).

### Statistical analysis

Statistical analyses were performed with GraphPad Prism version 6.0 software. Two tailed Student’s t-test was used to compare two independent groups. One-Way ANOVA was adopted to test the statistical differences between the means of two groups. Variables were described as means and standard deviations (SD). Statistical significance was indicated as follows: \**P* < 0.05; \*\**P* < 0.01.

## Results

### Retinoic acid secreted by activated hepatic stellate cells induces RARα nuclear translocation in liver progenitor cell line HepaRG cells

In acute liver failure, hepatic stellate cells (HSCs) are activated and secrete high levels of retinoic acid (RA) into the liver’s microenvironment where it can affect liver progenitor cells (LPCs). To investigate RA secretion by activated HSCs and its effect on LPCs, freshly isolated phHSCs and immortalized LX-2 HSCs were cultured for 96h or 72h respectively to determine RA concentrations and the expression of HSC activation markers. Activated phHSCs and LX-2 cells secreted more retinoic acid during culture as shown by absorbance measurements at 470 nm of conditioned cell culture supernatant **(Figure 1A)** and increased their HSC activation marker expression on mRNA level including that of *α-SMA*, *COL1α1*, *MMP2*, and *MMP9* **(Figure 1B**, **C**). Next, HepaRG cells were co-cultured with LX-2 cells for 72h. IF staining demonstrates that the RA receptor RARα was translocated into the nucleus of HepaRG cells as a result of LX-2-secreted RA **(Figure 1D)**. Further, atRA treatment also induced RARα nuclear translocation in HepaRG cells **(Figure 1E)**.

**Figure 1.**
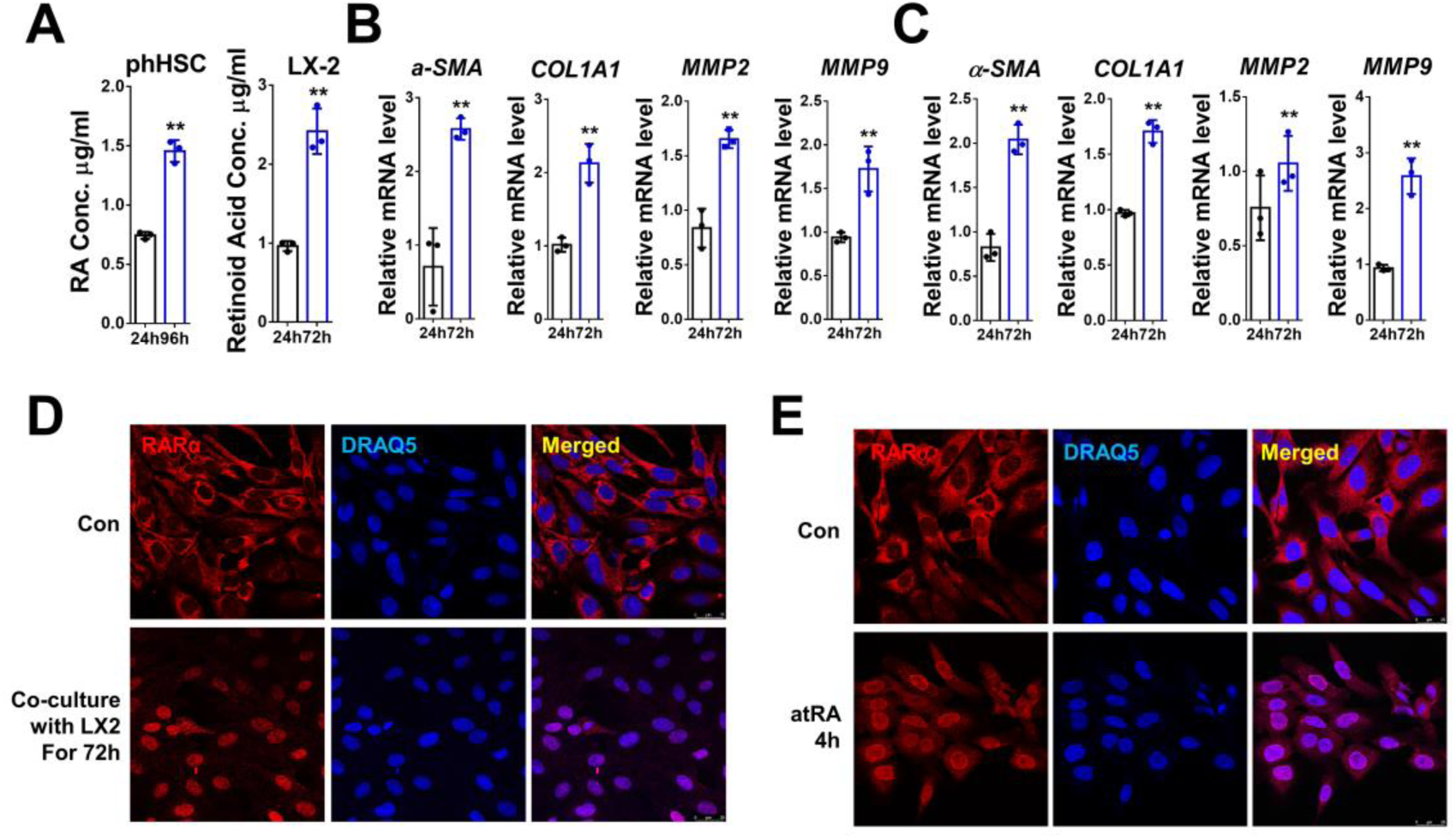
RA activates RARα signaling in HepaRG cells. **(A)** The concentration of retinoic acid (µg/ml) secreted by primary human hepatic stellate cells (phHSCs) or LX-2 cells by absorbance measurement at 352 nm. **(B, C)** Relative mRNA levels of *α-SMA, COL1A1, MMP2*, and *MMP9* were determined by qrtPCR in phHSCs and LX-2 cultured for 96h or 72h respectively. **(D, E)** Expression and location of RARα were detected by immunofluorescence staining in HepaRG cells co-cultured with/without LX-2 cells for 72h or treated with/without atRA for 4h. For qrtPCR, human PPIA was used as the endogenous control. Bars represent mean±SD. *p<0.05; **p<0.01. For IF, DRAQ5 was used to stain the nuclei. Scale bars, 25 µm. Images were chosen representatively from 3 independent experiments.

These data show that the RARα pathway in LPC is activated by HSC-secreted RA.

### Retinoic acid treatment significantly altered gene expression in HepaRG cells

To analyze the effect of RA on LPC, RNA sequencing (RNAseq) was used to analyze the transcriptomes of LPC, DMSO and atRA-treated HepaRG cells (n=3). Genes significantly affected by atRA treatment are shown in the heatmap **(Figure 2A)**. GO enrichment, enrichment score, and KEGG enrichment analysis showed that, in response to atRA treatment in HepaRG cells, the top-upregulated genes were strongly involved in, among others, cytokine-cytokine interaction, PI3K-AKT signaling, ECM-receptor interaction, and pathways related to cancer **(Figure 2B-D)**, while the most downregulated genes were related to metabolism, IL17 signaling pathway, insulin signaling pathway and cytokine-cytokine interaction **(Figure 2E-G)**. To investigate the effect of RA on LPCs in ALF, we mainly focused on the pathways crucial for LPC activation, including WNT and Hippo/YAP pathways, stem cell markers, metabolism, and survival-related signaling.

**Figure 2.**
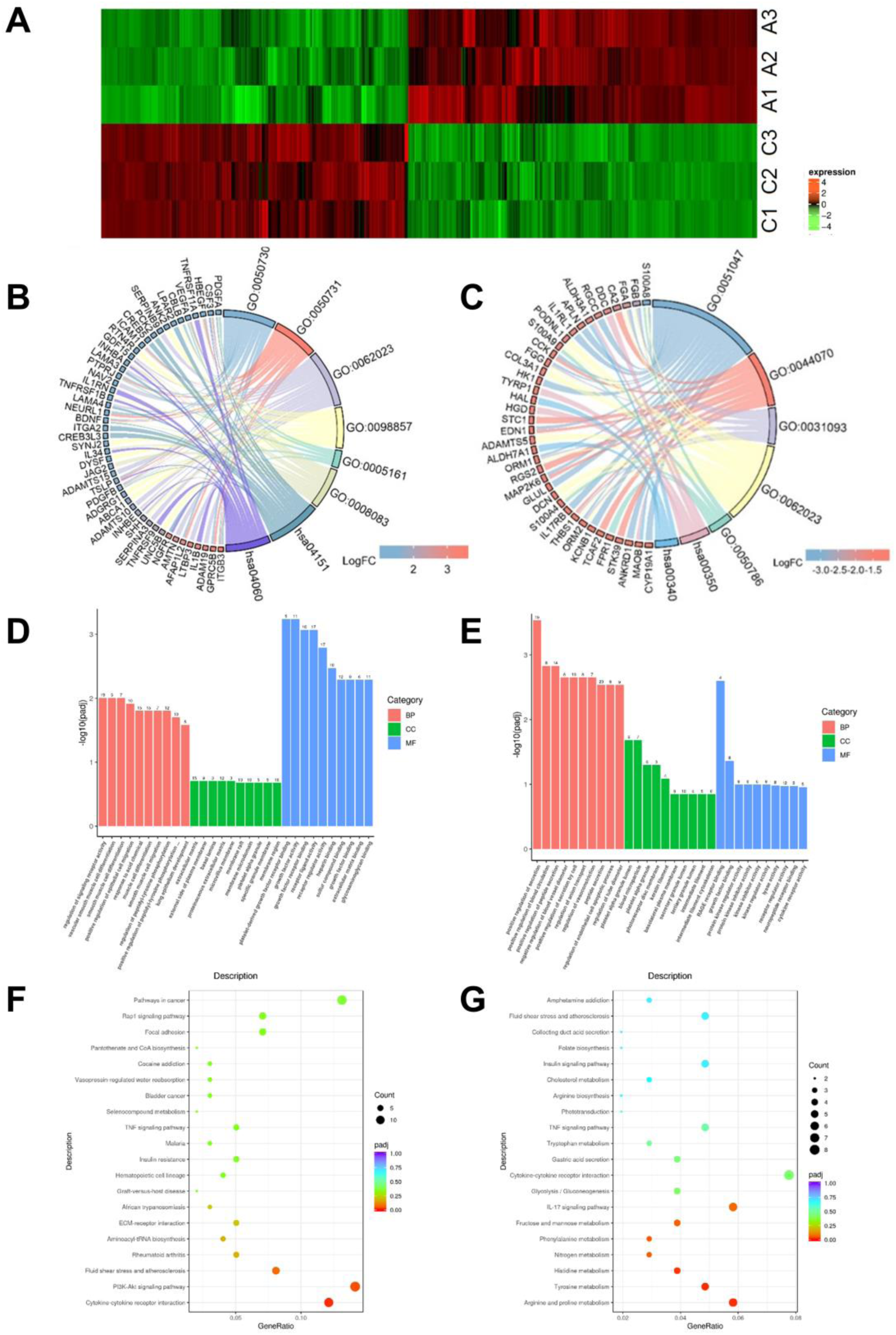
RNA-Seq analysis of HepaRG cells treated with control or atRA. **(A)** Heatmap of the upregulated and downregulated genes in response to atRA treatment in HepaRG cells. **(B, C)** Chord diagram, illustrating the main biological processes the downregulated or upregulated genes are involved in. **(D, E)** Enrichment score analysis of control vs atRA treatment. **(F, G)** GO enrichment analysis of control vs atRA treatment.

### Retinoic acid activates WNT signalling in HepaRG cells

According to RNAseq data, the expression of activators and members of the WNT signaling pathway were upregulated significantly upon atRA treatment **(Figure 3A)**. To verify WNT signaling activation, HepaRG cells were incubated with/without atRA for 24h following transfection with RARα siRNA or control siRNA. RARα knockdown efficiency by siRNA was verified by qrt PCR and WB **(Figure 3B)**. WNT signaling ligands WNT2b and WNT7B were induced by atRA treatment or co-culture with LX-2 cells corresponding to RNAseq data **(Figure 3C**, **D)**. In addition, atRA-induced WNT2B/7B expression was blunted, when RARα was knocked down **(Figure 3C**, **D)**. Further, active β-Catenin and total β-Catenin, the core proteins of the WNT signalling pathway, were both increased in response to atRA treatment for 4h, 8h, and 24h in HepaRG cells **(Figure 3E)** and depended on RARα expression **(Figure 3F)**. Co-culture with LX2 cells for 72h also promoted active β-Catenin and total β-Catenin expression **(Figure 3G)**. IF staining of active β-Catenin showed that its nuclear translocation was stimulated by atRA in HepaRG cells and disrupted by RARα knockdown **(Figure 3H)**. Total β-Catenin was localized on the cell membrane in control HepaRG cells in line due to its already established function in cell-cell adhesion. However, following atRA-treatment or co-culture with LX-2 cells, some of the β-Catenin was translocated into the nucleus as a central element of canonical WNT signaling **(Figure 3I**, **J)**. Taken together, atRA activated the WNT/β-Catenin pathway in HepaRG cells.

**Figure 3.**
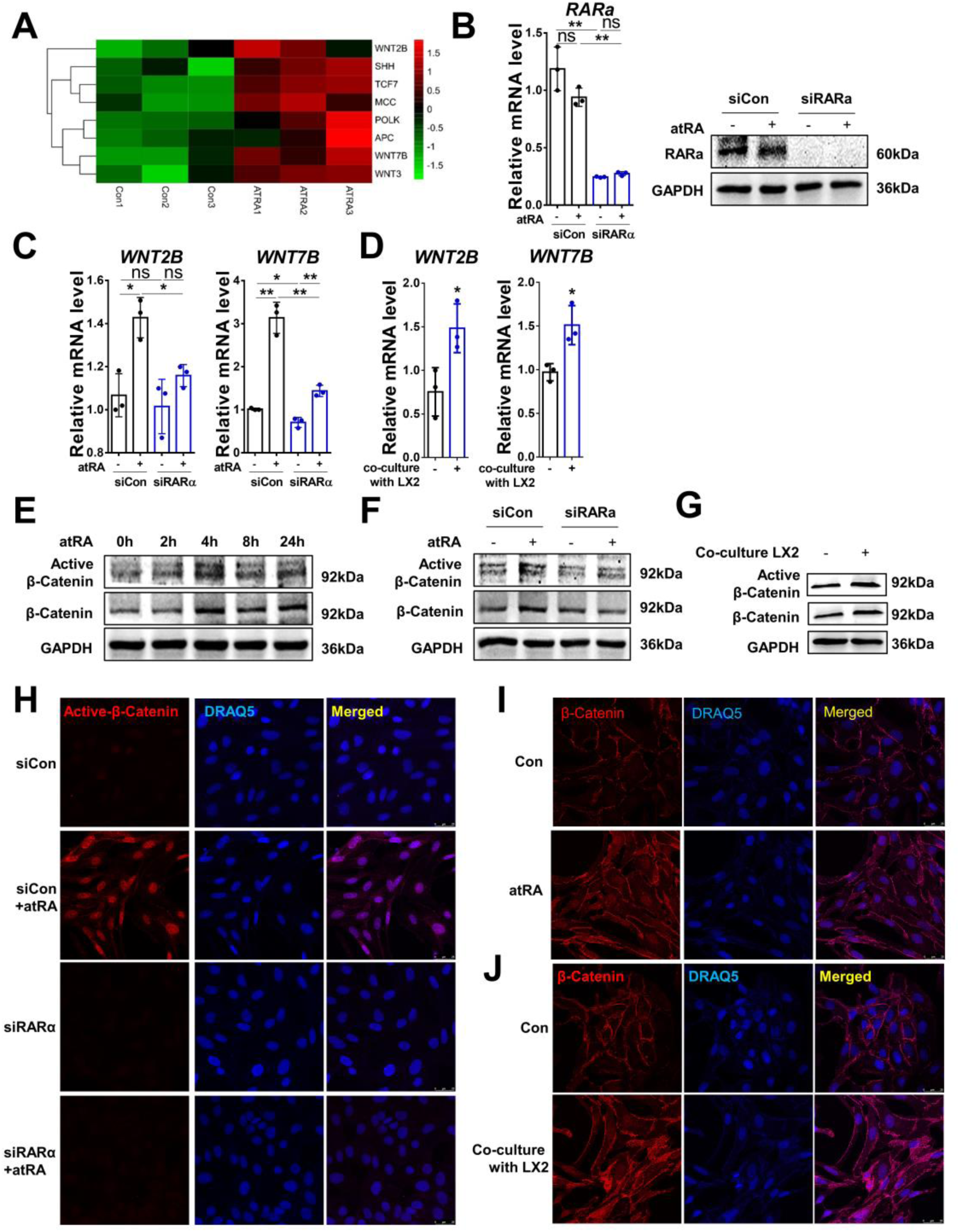
RA activates WNT signaling in HepaRG cells. **(A)** Heatmap showing the upregulated genes in WNT pathway. **(B)** Relative mRNA level and protein expression of RARα in HepaRG cells treated with/without atRA followed by transfection of control or RARα siRNA. **(C)** Relative mRNA levels of *WNT2B* and *WNT 7B* in HepaRG cells treated with/without atRA followed by transfection of control or RARα siRNA. **(D)** Relative mRNA levels of *WNT2B* and *WNT 7B* in HepaRG cells co-cultured with/without LX-2 cells for 72h. **(E)** Protein levels of active β-Catenin and total β-Catenin were determined by Western blotting in HepaRG cells treated with atRA for 0, 2, 4, 8, 24h. **(F)** Protein levels of active β-Catenin and total β-Catenin were determined by Western blotting in HepaRG cells treated with/without atRA followed by transfection of control or RARα siRNA. **(G)** Protein levels of active β-Catenin and total β-Catenin were determined by Western blotting in HepaRG cells co-cultured with/without LX-2 cells for 72h. **(H)** Location of active β-Catenin was detected by immunofluorescence staining in HepaRG cells treated with/without atRA followed by transfection of control or RARα siRNA. **(I, J)** Location of total β-Catenin was detected by immunofluorescence staining in HepaRG cells treated with/without atRA for 24h or co-cultured with/without LX-2 for 72h. For qrtPCR, human PPIA was used as the endogenous control. Bars represent mean±SD. *p<0.05; **p<0.01. For Western blotting, GAPDH was used as the loading control. For IF, DRAQ5 was used to stain the nuclei. Scale bars, 25 µm. Images were chosen representatively from 3 independent experiments.

### Retinoic acid enhanced the stemness of LPCs

Based on RNAseq, transcription factors involved in stemness were considerably increased following atRA treatment, including the well-studied stem cell markers, *SRY-box transcription factor 9* (*SOX9*), *alpha fetoprotein* (*AFP*), *CCAAT enhancer binding protein alpha* (*CEBPA*), *CCAAT enhancer binding protein gamma* (*CEBPG*), *GATA binding protein 6* (*GATA6*), *DNA polymerase kappa* (*POLK*), and *keratin 19* (*CK19*) **(Figure 4A)**. Among these transcription factors, *SOX9*, *AFP*, *CEBPA*, *GATA6*, *POLK* and *CK19* mRNA were increased in HepaRG cells after atRA incubation as verified by qrtPCR. Their expression was sustained by RARα expression **(Figure 4B)**. Co-culture with LX-2 cells also promoted *SOX9* and *CK19* expression on mRNA level **(Figure 4C)**. Western blotting confirmed that the stimulatory effect of atRA on SOX9 and AFP protein expression was inhibited by RARα knockdown **(Figure 4D)**. Co-culture with LX-2 cells also increased SOX9 expression on protein level in HepaRG cells, although AFP expression was not increased **(Figure 4E)**. IF staining displayed positive nuclear signal for SOX9 in HepaRG cells treated with atRA **(Figure 4F)** or co-cultured with LX-2 cells **(Figure 4G)**. atRA-induced SOX9 nuclear expression was blocked by transfection with RARα siRNA **(Figure 4F)**. Given that SOX9 is an important regulator of the transition between hepatic and biliary cell phenotypes during both liver development and regeneration, and that AFP is a well-known liver progenitor cell marker (7, 34, 35), we investigated SOX9 and AFP mRNA expressions in HepaRG cells cultured with growth medium (GM) or differentiation (LPC-to-hepatocyte differentiation) medium (DM) treated with/without atRA on days 2, 4, and 7. GM promoted SOX9 and AFP mRNA expression with atRA further enhancing this effect as opposed to DM. However, atRA incubation rescued the DM-mediated inhibitory outcome **(Figure 4H)**.

**Figure 4.**
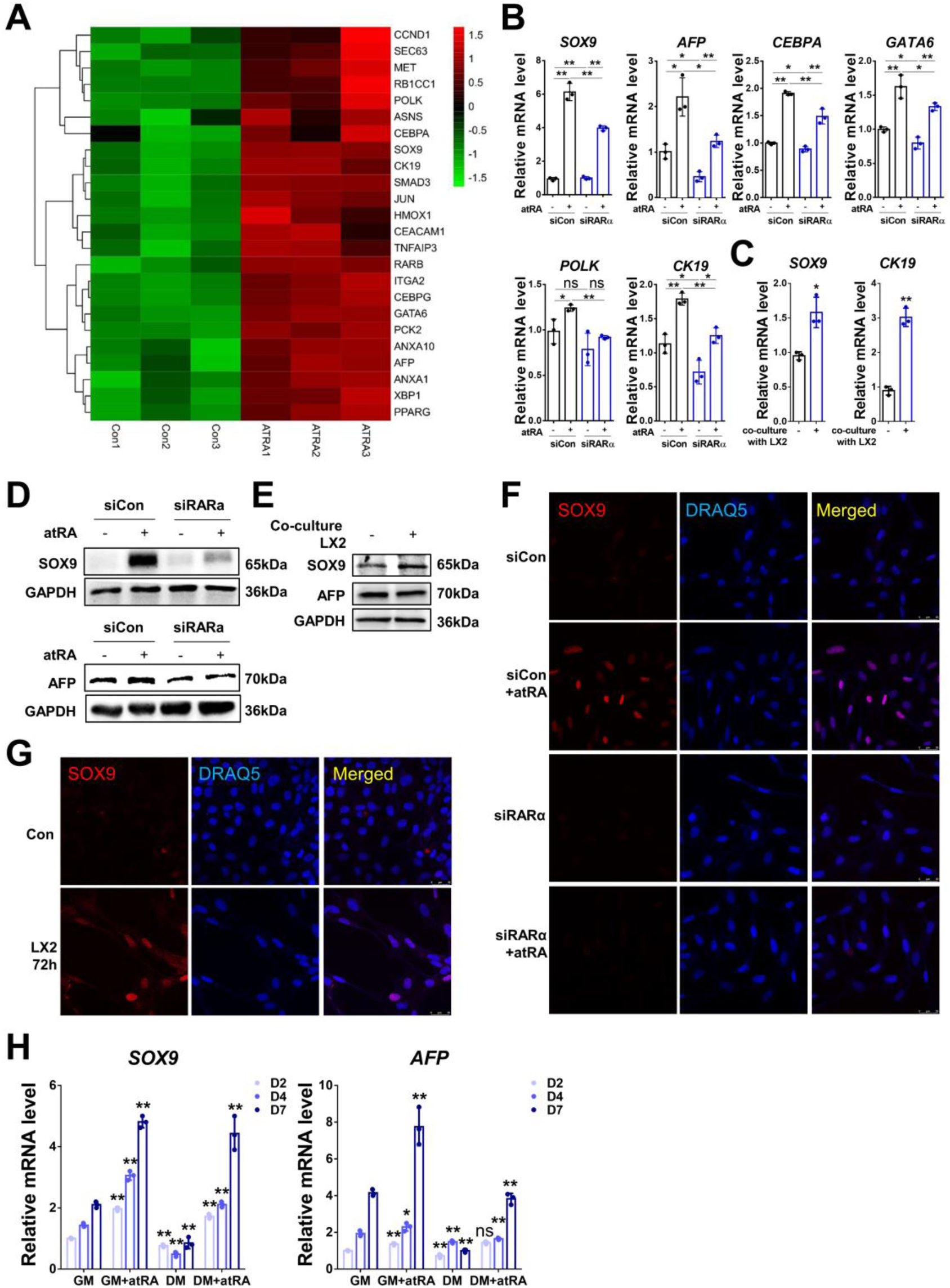
Retinoid acid enhanced the stemness of the LPCs. **(A)** Heatmap showing the upregulated genes involved in stemness maintenance. **(B)** Relative mRNA levels of *SOX9*, *AFP*, *CEBPA*, *GATA6*, *POLK*, and *CK19* in HepaRG cells treated with/without atRA followed by transfection of control or RARα siRNA. **(C)** Relative mRNA levels of *SOX9* and *CK19* in HepaRG cells co-cultured with/without LX-2 cells for 72h. **(D)** Protein levels of SOX9 and AFP were determined by Western blotting in HepaRG cells treated with/without atRA followed by transfection of control or RARα siRNA. **(E)** Protein levels of SOX9 and AFP were determined by Western blotting in HepaRG cells co-cultured with/without LX-2 cells for 72h. **(F)** Location of SOX9 was detected by immunofluorescence staining in HepaRG cells treated with/without atRA followed by transfection of control or RARα siRNA. **(G)** Location of SOX9 was detected by immunofluorescence staining in HepaRG cells co-cultured with/without LX-2 for 72h. **(H)** Relative mRNA levels of *SOX9* and *AFP* in HepaRG cells cultured in growth medium (GM) or differentiation medium (DM) incubated with/without atRA for 2, 4, and 7 days. For qrtPCR, human PPIA was used as the endogenous control. Bars represent mean±SD. *p<0.05; **p<0.01. For Western blotting, GAPDH was used as the loading control. For IF, DRAQ5 was used to stain the nuclei. Scale bars, 25 µm. Images were chosen representatively from 3 independent experiments.

These data suggested that atRA promotes the stemness of HepaRG cells by increasing the expression of stem cell markers.

### Retinoic acid promotes energy storage in the LPCs

The liver is an essential metabolic organ which governs the whole body’s energy metabolism (36). Under ALF conditions with most of the hepatocytes damaged or necrotic, LPCs are essential for the regulation of metabolism. Based on our RNA sequencing data, atRA promoted or inhibited the expression of several genes, depending on whether they are involved in lipogenesis or β-oxidation, respectively **(Figure 5A)**. We examined the mRNA expression of genes participating in lipogenesis upon atRA treatment. The qrtPCR data showed that atRA induced *perilipin 1* (*PLIN1*), *perilipin 5 (PLIN5*), *apolipoprotein A1* (*APOA1*), and *acyl-CoA synthetase long chain family member 5* (A*CSL5*) expressions on mRNA level, RARα-dependently **(Figure 5B)**. Unexpectedly, co-culture with LX-2 cells only increased *PLIN1* mRNA expression **(Figure 5C)**. This may be explained by the fact that, besides RA, activated HSCs also secrete TGF-β which inhibits lipogenesis and thus counteracts RA’s effect on fatty acid metabolism (37). Further, we tested the mRNA expression of genes involved in β-oxidation. qrtPCR validated atRA treatment or co-culture with LX-2 inhibiting *lipoprotein lipase* (*LPL*), *carnitine palmitoyltransferase 1B* (*CPT1B*), and *acetyl-CoA acyltransferase 1* (*ACAA1*) expression on mRNA level in HepaRG cells **(Figure 5D)**. Co-culture with LX-2 cells also reduced *LPL*, *CPT1B*, and *ACAA1* mRNA expression **(Figure 5E)**. Since CPT1B is the rate-controlling enzyme of the long-chain fatty acid β-oxidation pathway and ACAA1 is an enzyme involved in peroxisomal β-oxidation, these results implicate that atRA diminished fatty acid degradation in HepaRG cells. Notably, atRA incubation caused lipid droplet accumulation in HepaRG cells, which relied on RARα expression as well **(Figure 5F)**, although the inhibition of CPT1B and ACAA1 by atRA was independent of RARα **(figure 5D)**.

**Figure 5.**
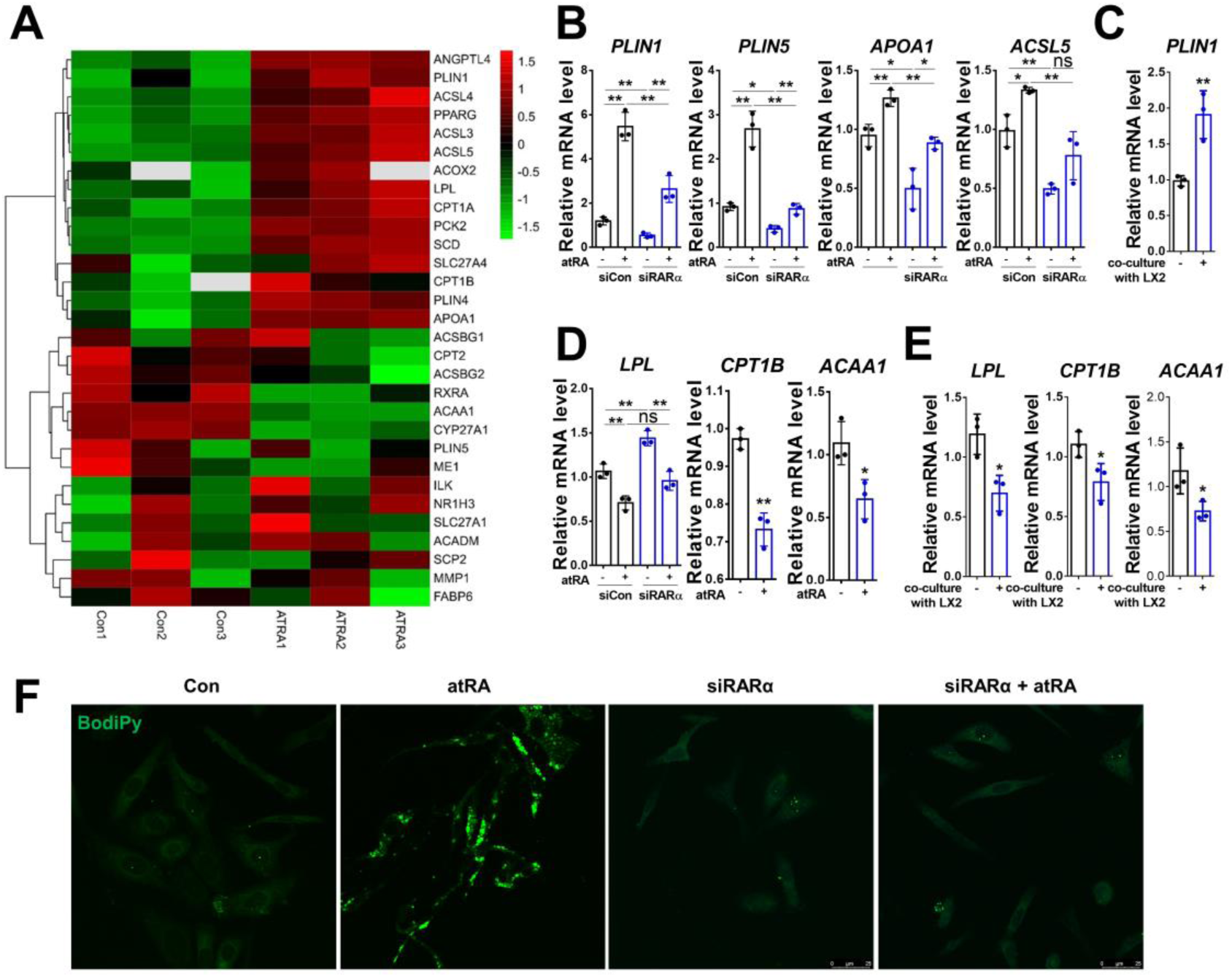
RA promotes energy storage in the LPCs. **(A)** Heatmap showing the downregulated and upregulated genes involved in metabolism upon atRA treatment in HepaRG cells. **(B)** Relative mRNA levels of *PLIN1*, *PLIN5*, *APOA1*, and *ACSL5* in HepaRG cells treated with/without atRA followed by transfection of control or RARα siRNA. **(C)** Relative mRNA levels of *PLIN1* in HepaRG cells co-cultured with/without LX-2 cells for 72h. **(D)** Relative mRNA levels of *LPL*, *CPT1B*, and *ACAA1* in HepaRG cells treated with/without atRA followed by transfection of control or RARα siRNA. **(E)** Relative mRNA levels of *LPL*, *CPT1B*, and *ACAA1* in HepaRG cells co-cultured with/without LX-2 cells for 72h. **(F)** Lipid droplet accumulation in HepaRG cells treated with/without siRARα and atRA was examined by BodiPy staining. DRAQ5 was used for the nuclear staining. For qrtPCR, human PPIA was used as the endogenous control. Bars represent mean±SD. *p<0.05; **p<0.01. For BODIPY staining, scale bars, 25 µm. Images were chosen representatively from 3 independent experiments.

These results illustrated that atRA promoted and inhibited gene expression with relation to fatty acid β-oxidation and lipogenesis respectively, most likely to promote energy storage in HepaRG cells. In the company of activated HSC, some of the impact of atRA on fatty acid metabolism is attenuated by TGF-β.

### Retinoid acid increases ABC transporters expression in HepaRG cells

Several of the superfamily of ATP binding cassette (ABC) transporters, were upregulated in response to atRA treatment in HepaRG cells, including, among others, *ABCG2*, *ABCA5*, *ABCA1*, *ABCC3*, and *ABCB4* **(Figure 6A)**. Verified by qrtPCR, atRA significantly induced the expression of *ABCC1*, *ABCC3*, and *ABCB4* on mRNA level in an RARα-dependent manner **(Figure 6B)**, which was further demonstrated by co-culture with LX-2 cells for 72h **(Figure 6C)**. Immunoblotting showed the increased expression of ABCC1/MRP1 on protein level in response to atRA stimulation, which was blocked by siRARα **(Figure 6D)**. Co-culture with LX-2 cells also increased MRP1 protein expression compared to control HepaRG cells **(Figure 6E)**. In addition, the increased membrane-bound expression of MRP1 in HepaRG cells triggered by atRA-RARα signaling was shown by IF staining **(Figure 6F)**.

**Figure 6.**
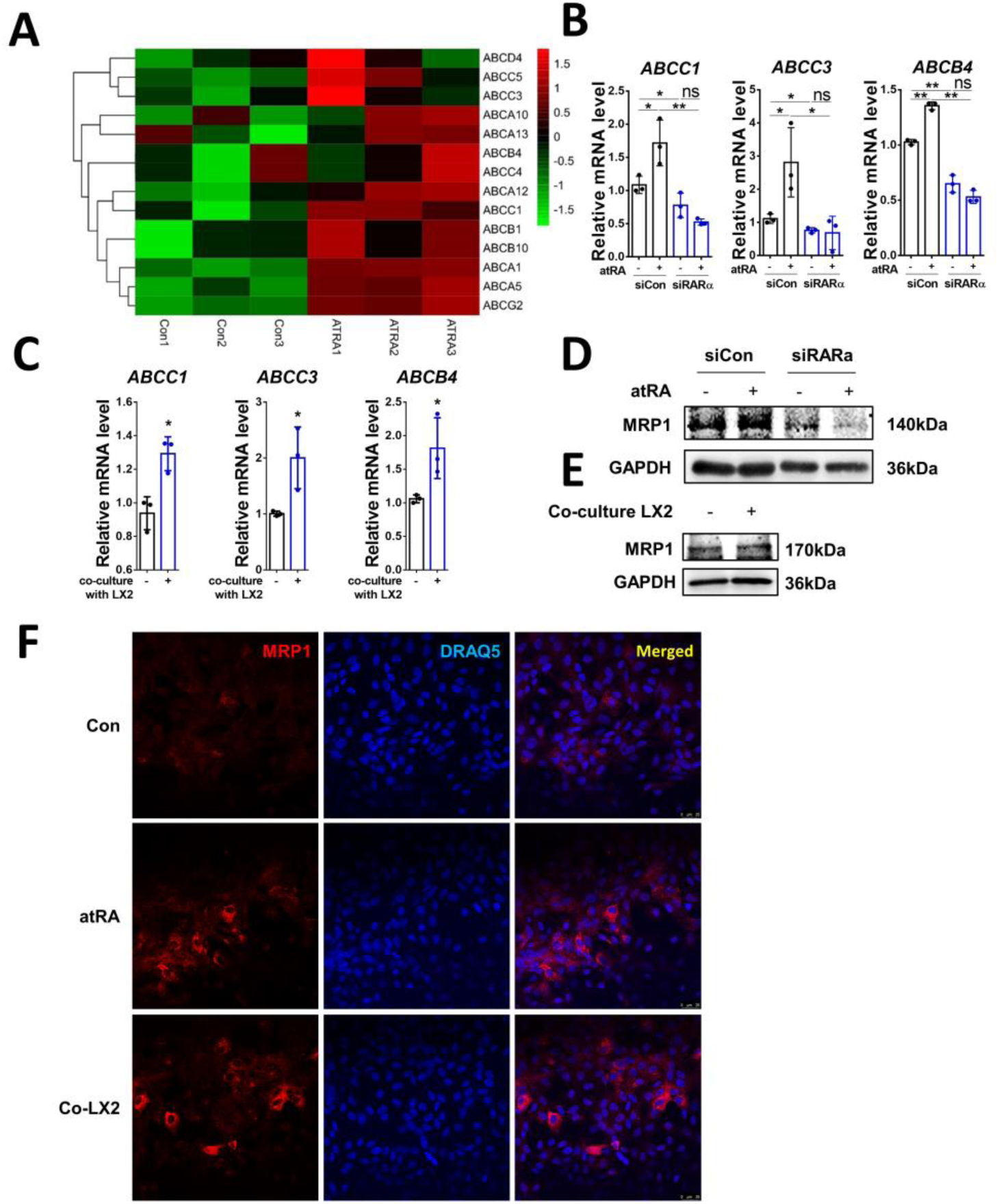
RA increases ABC proteins expression in HepaRG cells. **(A)** Heatmap showing the upregulated ABC-subfamily genes expression upon atRA treatment in HepaRG cells. **(B)** Relative mRNA levels of *ABCC1*, *ABCC3*, and *ABCB4* in HepaRG cells treated with/without atRA followed by transfection of control or RARα siRNA. **(C)** Relative mRNA levels of *ABCC1*, *ABCC3*, and *ABCB4* in HepaRG cells co-cultured with/without LX-2 cells for 72h. **(D)** Protein level of MRP1 was determined by Western blotting in HepaRG cells treated with/without atRA followed by transfection of control or RARα siRNA. **(E)** Protein level of MRP1 was determined by Western blotting in HepaRG cells co-cultured with/without LX-2 cells for 72h. **(F)** Cellular location of MRP1 was detected by immunofluorescence staining in HepaRG cells treated with/without atRA or co-culture with LX-2 cells for 72h. For qrtPCR, human PPIA was used as the endogenous control. Bars represent mean±SD. *p<0.05; **p<0.01. For Western blotting, GAPDH was used as the loading control. For IF, DRAQ5 was used to stain the nuclei. Scale bars, 25 µm. Images were chosen representatively from 3 independent experiments.

The proteins encoded by the ABC family play an important role in the transport of biliary compounds and especially, the intestinal excretion of organic anions and are involved in multi-drug resistance. atRA could promote these genes’ expression to generate a permissive niche for the survival of liver progenitor cells under ALF conditions.

### ALF patients with positive nuclear RARα in LPCs have a better prognosis

To corroborate our findings derived from mRNA sequencing analysis, we performed single-cell CyTOF analysis, and the schematic workflow is delineated in **Figure 7A**. The CyTOF-generated dataset underwent identification through single-cell proteomic analysis utilizing the t-distributed stochastic neighbor embedding (t-SNE) algorithm for comprehensive visualization of individual cells. The resulting t-SNE heatmap was portrayed as a bi-axial scatter plot representing a single cell. Each dot’s position on the plot was determined by the combined expression of all proteins within that specific cell. The t-SNE heatmap of β-Catenin, SOX9, and AFP indicated that while there was no significant difference in the protein expression patterns between the two cell groups, the red dots denoting high expression levels were found in atRA-treated HepaRG cells **(Figure 7B)**. The corresponding protein expression values for each sample were also depicted in the histograms **(Figure 7C)**. To gain deeper insights into the mechanism underlying atRA/RARα-induced upregulation of SOX9 and AFP, a chromatin immunoprecipitation (ChIP) assay was conducted. The results revealed a significant induction of RARα binding to the gene promoter regions of *CTNNB1* (−118 ∼ -267) with predicted binding sites AGGCCGCGCAGCGGGCA and *SOX9* (−5 ∼ -108) with predicted binding sites GGGTGGCTCTAAGGTGA following incubation with atRA **(Figure 7D**, **E)**. Given the indications from our findings that atRA promoted LPC activation, characterized by increased cell proliferation, we conducted a MTT assay. The results demonstrated that incubation with atRA for 72 hours significantly enhanced HepaRG cell viability **(Figure 7F)**, potentially through the upregulation of *CCND1*, *Ki67*, and *CDK6* **(Figure 7G)**.

**Figure 7.**
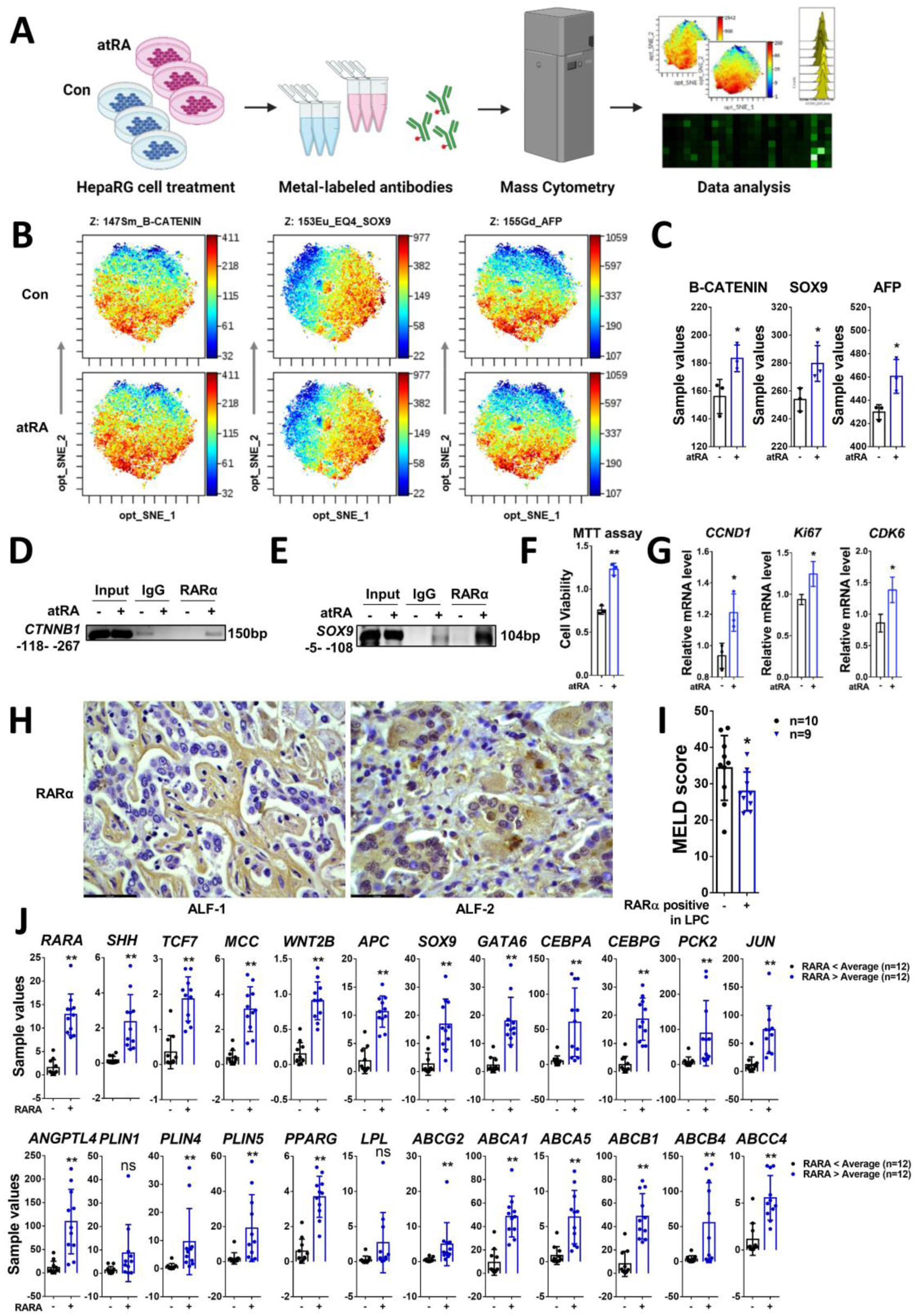
ALF patients with positive nuclear RARα in LPCs have a better prognosis. **(A)** CyTOF analysis of atRA effects on HepaRG cells (Created with BioRender.com). **(B, C)** t-SNE heatmap and sample values of protein expression among the control and atRA treatment group. **(D, E)** ChIP assay of RARα binding to the *CTNNB1*, and *SOX9* gene promoter in HepaRG cells upon control or atRA treatment with the PCR amplified products were shown on a 2% agarose gel. Fragment “-118bp ∼ -267bp’’ or “-5bp ∼ -108bp’’ was relative to the transcription start site of *CTNNB1* or *SOX9* gene respectively. Rabbit IgG-bound chromatin served as a negative control. **(F)** MTT assay of HepaRG cells with control or atRA incubation for 72h. **(G)** Relative mRNA levels of *CCND1*, *Ki67*, and *CDK6* in HepaRG cells treated with/without atRA. **(H)** IHC staining of RARα in the liver tissue of 21 ALF patients. **(I)** MELD score of all ALF patients with or without RARα nuclear expression. **(J)** mRNA expression of *RARA* and the target genes in liver tissue of 24 patients with ALF, extracted from GEO DataSets GSE164397. Bars represent mean±SD. *p<0.05; **p<0.01. Scale bars, 43.5 µm. Images were chosen representatively from IHC staining.

Next, we performed IHC staining of RARα in the liver tissue of patients with ALF. 19 ALF patients were divided into two cohorts in accordance with RARα nuclear staining, staining either positively or negatively in LPCs for RARα and the MELD score of every patient was calculated. 10 patients showed negative RARα nuclear staining while 9 patients stained positive **(Figure 7H**). ALF patients that maintained nuclear RARα expression had a better MELD score **(Figure 7I)**, implying that RARα signaling is correlated with a better prognosis in ALF patients. Lastly, to verify the correlation of RARα activation and our target signaling pathways (Figure 3-6) in patients suffering from acute liver failure, we extracted the related genes from the Gene Expression Omnibus (GEO) dataset (GSE164397) that comprises mRNA expression data of liver tissue from pediatric ALF patients. 24 ALF patients were divided into *RARA*-(n=12) or *RARA*+ (n=12) groups according to the expression value of *RARA* being above or below its average. In *RARA*+ ALF patients, WNT signaling (*SHH*, *TCF7*, *MCC*, *WNT2B*, *APC*), stemness markers (*SOX9*, *GATA6*, *CEBPA*, *CEBPG*, *PCK2*, *JUN*), lipogenesis (*ANGPTL4*, *PLIN1*, *PLIN4*, *PLIN5*, PPARG), and ABC transporters (*ABCG2*, *ABCA1*, *ABCA5*, *ABCB1*, *ABCB4*, *ABCC4*) were significantly upregulated **(Figure 7J)**, which was in accord with our findings in HepaRG cells.

## Discussion

The liver is historically famous for its enormous regenerative capacity under the most adverse conditions. There are two main ways of liver regeneration, hepatocytes- and/or LPC-mediated regeneration (38, 39). The primary mode of regeneration is driven by already differentiated hepatocytes undergoing proliferation; the second mode requires differentiation of hepatic stem cells LPCs into hepatocytes. The latter mode plays a crucial role when the majority of hepatocytes no longer perform effectively, for instance in the setting of ALF (5, 38, 39). The initiation of LPC-mediated regeneration requires LPC activation, which corresponds histologically to the so-called ductular reaction (5).

To date, the signals which are needed within the microenvironment of ALF livers to initiate LPC activation and the subsequent process of regeneration remain largely unknown. In this study, we revealed that retinoic acid/RARα signaling may be responsible for LPC activation by enhancing their stemness, increasing energy storage, and increasing drug resistance **(Figure 8)**, all of which are significant for subsequent liver regeneration and correlate with a better prognosis for ALF patients.

**Figure 8.**
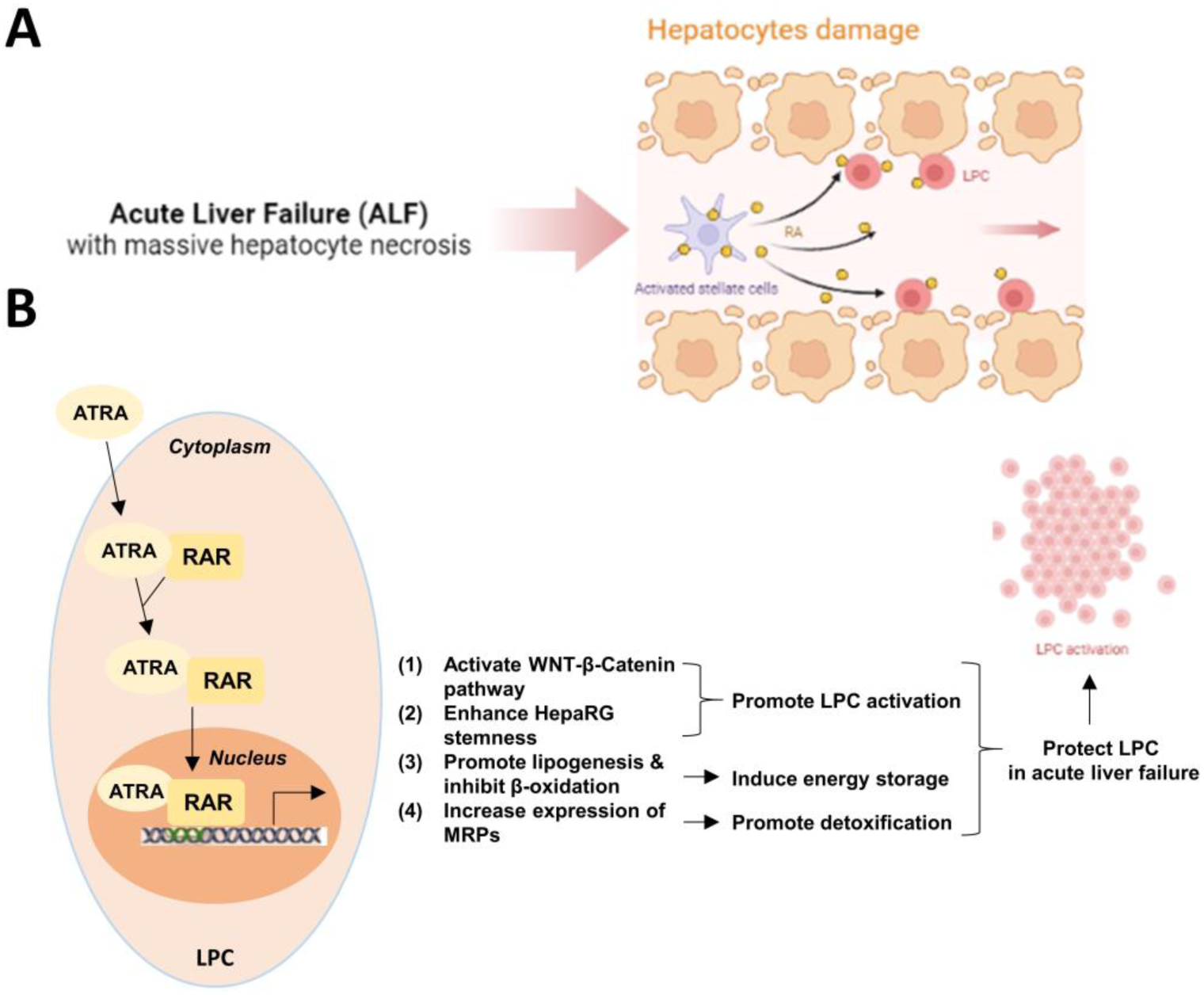
Schematic Diagram of RA’s role in regulating LPC activation under ALF conditions. **(A)** In ALF, activated HSCs secrete high amounts of RA. Since most of the hepatocytes are damaged or deceased, LPCs are the main cells capable of conveying RA signaling. **(B)** RA translocates with RARα into the nuclei in LPCs to induce the transcription of retinoid-responsive genes, including members of the WNT/β-Catenin pathway, regulating cellular stemness, metabolic regulation, and multi-drug resistance in order to enhance the stemness, energy storage, and detoxification in LPCs, thus enabling the LPCs to survive and drive liver regeneration under ALF conditions. (Created with BioRender.com.)

To investigate the stemness of HepaRG cells treated with atRA, we analysed RNAseq data and verified them by qrtPCR, revealing that WNT signaling and LPC makers including SOX9, AFP, and CK7 are significantly upregulated. The WNT/β-Catenin pathway is a highly conserved and tightly controlled molecular mechanism that regulates hepatobiliary development and cell fate during embryogenesis, as well as liver homeostasis and repair in adulthood (40, 41). Besides, WNT signaling is essential during liver regeneration as it is key for the activation of oval cells, which are liver progenitor cells (42). Additionally, WNT seems to play a role in LPC expansion and differentiation (43). However, aberrant activation of the WNT/β-Catenin pathway entails a high risk of hepatocellular carcinoma (HCC) development (44–46), implying that atRA/RAR signaling needs to be targeted precisely to specific stages during ALF. It will be essential to target the initiation stage of ALF in which the LPCs need to be activated rather than a later stage, when the LPCs differentiate into hepatocytes, as the latter might promote HCC development. Our results also revealed that SOX9, AFP, and CK19 are induced significantly by atRA treatment. These are all generally accepted LPC markers (7). However, the underlying mechanisms of their regulation as well as the extent of the involvement of WNT signaling and its own regulation by atRA are not fully understood. Whether atRA affects these pathways through RARα binding to the respective promoters directly or through a crosstalk between SOX9 and WNT/β-Catenin still warrants further investigation. Previous studies found that as β-Catenin lacks the intrinsic ability to bind to DNA, its interaction with target genes relies on the recruitment of co-activators and other transcription factors, such as hypoxia-inducible factor 1α (HIF1α) (47), forkhead box protein O (FOXO) (48), and SOX family transcription factors (49). Combining these findings with our own results, we suppose that atRA/RARα may activate WNT/β-Catenin signaling to induce SOX9 upregulation, leading to increased LPC marker expression, e.g., AFP, as we found that knockdown of SOX9 abolished atRA-induced AFP expression (Figure S1).

The liver is an important metabolic organ and plays an essential role in the body’s vital energy metabolism (36). Ensuring a sufficient energy supply for glucose-dependent organs and cells, such as the brain, heart, and in the liver LPCs, is crucial for the body’s resilience to a systemic illness such as ALF(50). Therefore, we examined the regulation of metabolic processes by atRA in HepaRG cells. We demonstrated that atRA inhibits fatty acid β-oxidation while at the same time promoting lipogenesis in LPCs, opposing its function in hepatocytes (16, 51), which we assume is for the reason that boosting their energy storage is essential for LPC survival under ALF conditions. Nonetheless, the detailed mechanisms need further study. We also found that ABC-family transporters were significantly induced by atRA. Besides their role in cholesterol and bile acid transport, they are also involved in detoxification and multi-drug resistance, which are thought to be protective for LPCs (52). Several studies found that hepatic expression of ABCC2, ABCC3, ABCC4, and ABCB11 are regulated by atRA-RAR/RXR in hepatocytes (53–55), but there are not sufficient studies in LPCs yet. Based on our analyses, in LPCs, ABC transporters can be upregulated by atRA, partially dependent on RARα, with the underlying mechanisms also warranting further investigation.

We show that ALF patients maintaining nuclear expression of RARα in LPCs have better MELD scores which translates to better survival. However, other clinical outcomes of our patients should also be analysed in future to confirm our correlation, such as the likelihood of sepsis, ascites, bacterial infections, renal failure, encephalopathy during ALF. In conclusion, our study assessed the effects of RA secreted by activated HSCs on LPCs and implies that the RA-RAR pathway may serve as a potential target in order to activate LPCs, especially at the early stage of ALF and promote liver regeneration, which could potentially lead to better treatment options for people suffering from ALF.

## Acknowledgement

We acknowledge the support of the LIMa Live Cell Imaging at Microscopy Core Facility Platform Mannheim (CFPM).

## Author’s contributions

Conception, design and hypothesis: Shanshan Wang

*In vitro* experiments: Sai Wang, Frederik Link, Stefan Munker, Shanshan Wang, Wenjing Wang, Rilu Feng, Yujia Li, and Ye Yao

Patients sampling and pathological experiments: Sai Wang, Frederik Link, Hui Liu, Chen Shao, Huiguo Ding, and Honglei Weng

Bioinformatics analysis: Sai Wang, Frederik Link, and Shanshan Wang Drafting the article: Sai Wang, Frederik Link, and Shanshan Wang

Data discussion, reviewing and editing the article critically: Sai Wang, Frederik Link, Shanshan Wang, Roman Liebe, Honglei Weng, Steven Dooley, Huiguo Ding, and Matthias P. A. Ebert

## Abbreviations

ABC: ATP-binding cassette
ACAA1: acetyl-CoA acyltransferase 1
ACSL5: acyl-CoA synthetase long chain family member 5
AFP: alpha fetoprotein
ALF: acute liver failure
APOA1: apolipoprotein A1
α-SMA: Alpha-smooth muscle actin
ATRA: all-trans retinoid acid
CEBP: CCAAT enhancer binding protein
CK7/19: Cytokeratin 7/19
COL1A1: Collagen, type I, alpha 1
CPT1B: palmitoyltransferase 1B
GATA6: GATA binding protein 6
HSC: hepatic stellate cell
IF: immunofluorescence
IHC: immunohistochemistry
INR: international normalized ratio
LPC: liver progenitor cell
LPL: lipoprotein lipase
LTx: liver transplantation
MELD: Model for End-Stage Liver Disease
MHN: Massive hepatic necrosis
MMP2/9: matrix metalloproteinase 2/9
PLIN1/5: perilipin 1/5
phHSC: primary human hepatic stellate cell
POLK: DNA polymerase kappa
RAR: retinoic acid receptor
RXR: retinoid X receptor
SOX9: SRY-box transcription factor 9
TBIL: total bilirubin
TGF-β1: transforming growth factor-β1
ULN: upper limit of normal

