## Supplementary material for "Retinoic acid generates a beneficial microenvironment for liver progenitor cell activation in acute liver failure"

**Table S1. Primers for PCR**

| Primer | Forward | Reverse |
|---|---|---|
| RARα | AAGCCCGAGTGCTCTGAGA | TTCGTAGTGTATTTGCCCAGC |
| α-SMA | GTGTTGCCCCTGAAGAGCAT | GCTGGGACATTGAAAGTCTCA |
| COL1A1 | GAGGGCCAAGACGAAGACATC | CAGATCACGTCATCGCACAAC |
| MMP-2 | TACAGGATCATTGGCTACACACC | GGTCACATCGCTCCAGACT |
| MMP-9 | TGTACCGCTATGGTTACACTCG | GGCAGGGACAGTTGCTTCT |
| PPIA | AGGGTTCCTGCTTTCACAGA | CAGGACCCGTATGCTTTAGG |
| WNT2B | GTTACCCAGACATCATGCGTT | GGGTGGTACAGTTCCAGCG |
| WNT7B | GAAGCAGGGCTACTACAACCA | CGGCCTCATTGTTATGCAGGT |
| SOX9 | AGCGAACGCACATCAAGAC | CTGTAGGCGATCTGTTGGGG |
| AFP | CTTTGGGCTGCTCGCTATGA | GCATGTTGATTTAACAAGCTGCT |
| CEBPA | GCGGCGGCGGCGACTTT | GGTAGCCGGCGGCCGCGCA |
| GATA6 | CTCAGTTCCTACGCTTCGCAT | GTTGGCACAGGACAATCCAAG |
| POLK | ACTTTGACAAATACCGAGCTGTG | GGAGAGATGGATCGTTCATGC |
| CK19 | ACCAAGTTTGAGACGGAACAG | CCCTCAGCGTACTGATTTCCT |
| PLIN1 | TGGGTGGTGTGGCACATAC | CCTCCCCTTGGTTGAGGAGA |
| PLIN5 | AAGGCCCTGAAGTGGGTTC | GCATGTGGTCTATCAGCTCCA |
| APOA1 | CCCTGGGATCGAGTGAAGGA | CTGGGACACATAGTCTCTGCC |
| ACSL5 | TGGCTATCTTACAAACAGGTGTC | TCCACTCTGGCCTATTCTGAG |
| LPL | ACAAGAGAGAACCAGACTCCAA | GCGGACACTGGGTAATGCT |
| CPT1B | CATGTATCGCCGTAAACTGGAC | TGGTAGGAGCACATAGGCACT |
| ACAA1 | ATGTGGCTGAGCGGTTTGG | GGCGGATACCCTCATCCTG |
| ABCC1 | GTCGGGGCATATTCCTGGC | CTGAAGACTGAACTCCCTTCCT |
| ABCC3 | TGGGGTGAAGTTTCGTACTGG | CACGTTTGACTGAGTTGGTGATA |
| ABCB4 | ATAGCTCACGGATCAGGTCTC | GGATTTAGCAGCGACAAGGAAA |
| ChIP-CTNNB1 | CCCCGATGCAGACCACAG | GGTGGAAAGTAGTCCCCGC |
| ChIP-SOX9 | ATCGGGTCCAATCAGCTGCC | CCCAGCCCAGGGTCTCTTTA |

*Supplementary figure and figure legend*

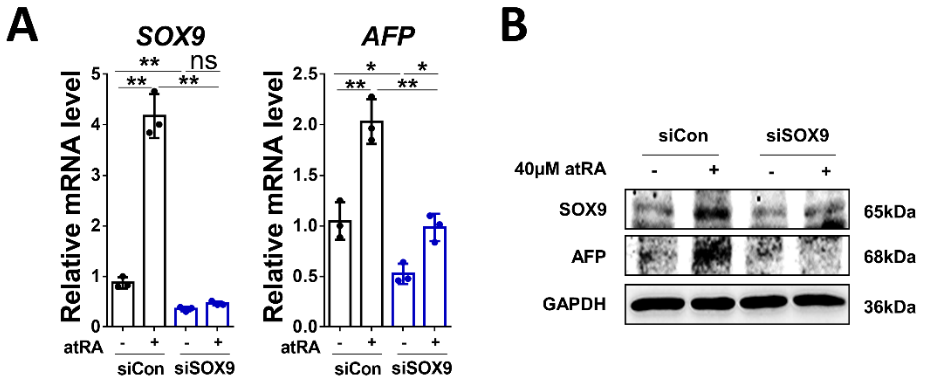


**Figure S1. SOX9 knockdown inhibited atRA-induced AFP expression. (A, B)** Relative mRNA and protein levels of SOX9 and AFP in HepaRG cells treated with control or atRA followed by RARα knockdown.

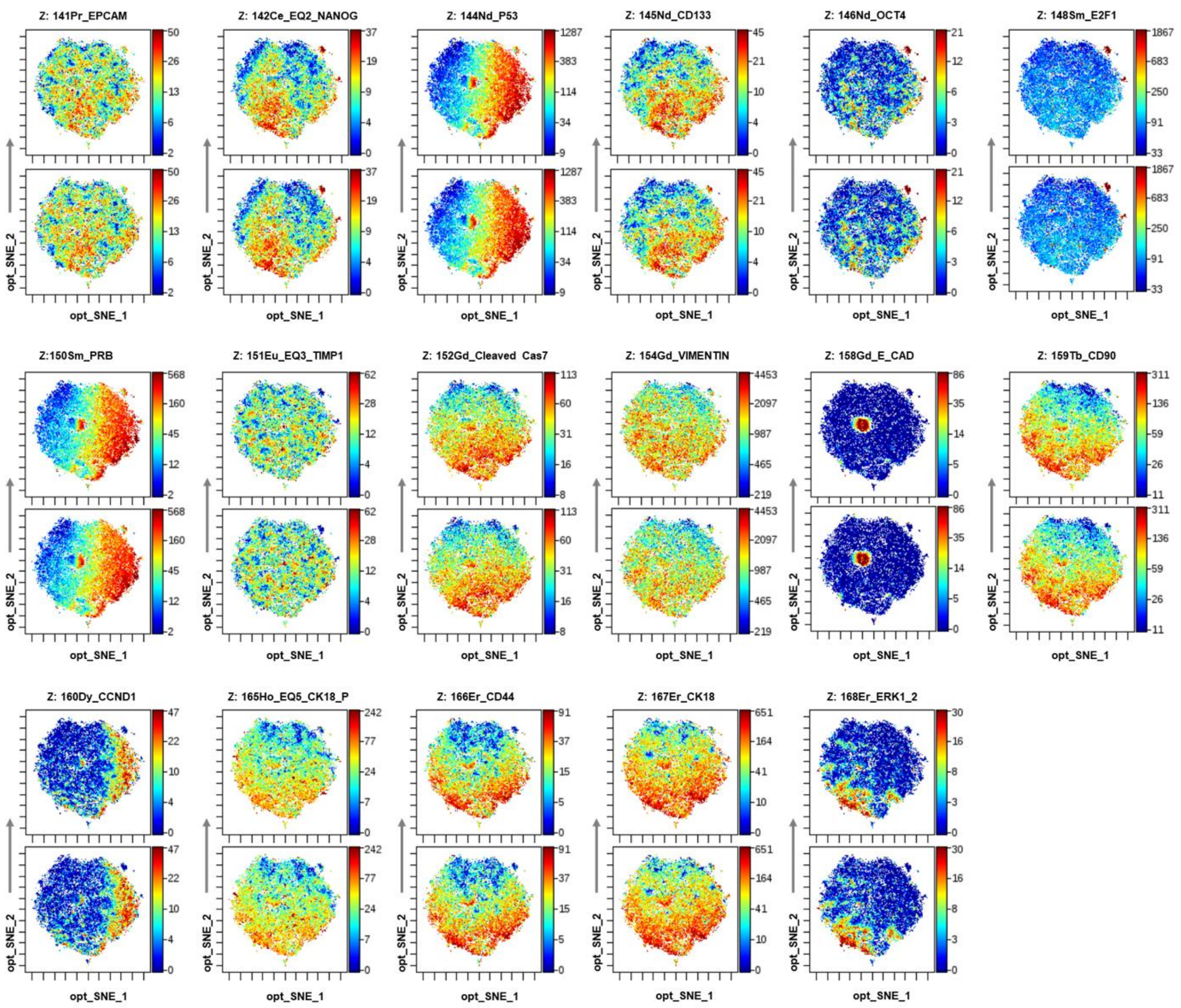


**Figure S2. t-SNE heatmap of protein markers expression among the control and atRA treatment group.**
